# Personalization of Closed-Chain Shoulder Models Yields High Kinematic Accuracy for Multiple Motions

**DOI:** 10.1101/2024.12.19.629415

**Authors:** Claire V. Hammond, Heath B. Henninger, Benjamin J. Fregly, Jonathan A. Gustafson

## Abstract

The shoulder joint complex is prone to musculoskeletal issues, such as rotator cuff-related pain, which affect two-thirds of adults and often result in suboptimal treatment outcomes. Current musculoskeletal models used to understand shoulder biomechanics are limited by challenges in personalization, inaccuracies in predicting joint and muscle loads, and an inability to simulate anatomically accurate motions. To address these deficiencies, we developed a novel, personalized modeling framework capable of calibrating subject-specific joint centers and functional axes for the shoulder complex. Leveraging in vivo biplane fluoroscopy data and the recent Joint Model Personalization Tool from the Neuromusculoskeletal Modeling Pipeline, we optimized joint parameters and body scale factors for shoulder models with varying degrees of freedom (DOFs). We initially created and tested open-chain scapula-only models (3DOF, 4DOF, and 5DOF) and found that increasing DOFs improved accuracy, with the 5 DOF model yielding the lowest marker distance errors (average = 0.8 mm, maximum = 5.2 mm) as compared to biplane fluorscopy data of the scapula across eight movement trials. We subsequently created closed-chain shoulder models incorporating scapula, clavicle, and humerus bodies. We found closed-chain shoulder models with 5 DOFs for the scapula achieved the highest accuracy (average = 0.9 mm, maximum = 5.7 mm) and showed consistent performance across subjects (n=3) in leave-one-out cross-validation tests (average marker distance errors = 1.0–1.4 mm). This framework minimizes errors in joint kinematics and provides a foundation for future models incorporating personalized musculature and advanced simulations, enhancing its potential clinical utility for rehabilitation and surgical planning.

## 1 Introduction

The shoulder joint complex allows for the largest range of motion in the human body. However, this high mobility comes at the cost of low joint stability that is controlled actively by the surrounding musculature of the rotator cuff and passively by the soft tissue (e.g., capsule, labrum, ligaments). The delicate balance of both mobility and stability in the shoulder complex helps explain why it is one of the most frequent sites of musculoskeletal complaints, with 50% of all initial episodes of shoulder pain becoming chronic (Luime et al., 2004; GBD 2017 Disease and Injury Incidence and Prevalence Collaborators, 2018). Acute to chronic shoulder injuries, like rotator cuff-related pain, affect two in three adults in their lifetime (Luime et al., 2004; Gray et al., 2016). While rotator cuff tears are the most common injury in the shoulder—followed by shoulder fracture and dislocation, the pathophysiology of shoulder injuries is complex due to the interaction of tendon, bone, and muscle that span both the glenohumeral and scapulothoracic joints (Bedi et al., 2024). Both surgical and non-surgical treatments have had varying success, with up to 50% of people remaining symptomatic 18 months after presentation of symptoms (Croft et al., 1996). The progression of shoulder-related pain to glenohumeral arthritis and/or rotator cuff arthropathy can vary significantly across individuals and underscores the lack of knowledge about how different treatments interact with patient-specific shoulder anatomy to determine post-treatment shoulder function (Croft et al., 1996).

Musculoskeletal computer modeling is a growing approach for investigating how different treatments may interact with a patient’s shoulder anatomy. Researchers have used musculoskeletal shoulder models to understand the effects of both rotator cuff repair and joint arthroplasty on muscle moment arms and joint reaction forces across the shoulder (Gutiérrez et al., 2008; Roche et al., 2009; Walker et al., 2016; Elwell et al., 2018; Leschinger et al., 2019; Huish et al., 2021). A major weakness of prior work is how these studies modeled the kinematic structure of the shoulder, and specifically of the scapulothoracic joint. The Delft Shoulder model, originally proposed in 1994 by van der Helm, included three joints with the scapulothoracic joint constrained to move along a gliding plane (van der Helm, 1994; Nikooyan et al., 2011). Wu et al. employed a ball-and-socket joint to model scapulothoracic motion, prioritizing accurate muscle paths and moment arms from cadaveric studies (Wu et al., 2016). Saul et. al created a shoulder model that utilized a thorax-fixed regression model to determine clavicle and scapula kinematics from static humeral poses—a common approach for modeling scapulothoracic motion (Holzbaur et al., 2005). A recent shoulder model created by Seth et. al utilized a coordinate mobilizer to add a degree of freedom to the scapula to describe winging, and the achievable motions were compared to measured bone-pin shoulder kinematic data for a single subject (Seth et al., 2016).

These existing shoulder models possess three important deficiencies. First, these models are difficult to personalize due to their use of regression-based kinematics and/or incompatibility with existing personalization tools. Second, these models either assume an open kinematic chain—which leads to less accurate muscle and joint load predictions—or contain artificially added degrees of freedom (DOFs) that allow for non-anatomical shoulder motion. Third, these models cannot be used for predicting novel motions because they lack consistent, closed kinematic chain structures. For a shoulder model to be clinically applicable, it must be able to reproduce in vivo shoulder motion with good accuracy. Thus, the deficiencies above limit the clinical applicability of existing shoulder models, particularly for rehabilitation or surgical planning purposes.

To address these deficiencies, we propose a novel personalized modeling framework that can calibrate subject-specific joint centers and functional axes for the shoulder complex. We address two past challenges that have limited the ability of the modeling community to create personalized shoulder kinematic models. The first challenge has been obtaining highly accurate in vivo kinematic data for multiple shoulder motion tasks. Such data have recently been collected from healthy subjects using bi-plane video fluoroscopy, and these data have been made freely available to the research community (Aliaj et al., 2021; Kolz et al., 2021; Henninger, n.d.). The second challenge has been developing a kinematic model personalization process that makes it easy to calibrate a shoulder model’s joint parameters (i.e., joint locations and orientations of joints in the body segments) such that the calibrated model with joint constraints can closely reproduce the unconstrained six DOF kinematics of the scapula and humerus. The Joint Model Personalization (JMP) Tool available within the recently released Neuromusculoskeletal Modeling Pipeline software provides such a process (Hammond et al., 2024). Using the JMP Tool with in-vivo kinematic data available from three subjects, we developed personalized shoulder models with consistent, closed kinematic chain structures that closely reproduced in vivo scapulothoracic and glenohumeral kinematics.

## 2 Model & Methods

### 2.1 Overview

We developed three personalized shoulder kinematic models using subject-specific bony anatomy and kinematic data from an open-source repository (Aliaj et al., 2021; Kolz et al., 2021; Henninger, n.d.). The models were formulated with varying degrees of freedom (DOF) to evaluate their accuracy and generalizability. As one of our primary goals was to develop a modeling approach to accurately track the scapula, we initially created scapula-only kinematic models with three, four, and five DOFs (Figure 1). Using knowledge gained from the scapula-only model, we subsequently created a closed-chain kinematic model of the shoulder incorporating scapula, clavicle, and humerus bodies. We evaluated the accuracy of the kinematic models by comparing marker positions produced by each model with synthetic marker positions generated from subject-specific geometric models and biplane fluoroscopy data. Using the Joint Model Personalization (JMP) Tool available within the Neuromusculoskeletal Modeling Pipeline software (Hammond et al., 2024), we optimized joint parameters and body scale factors to minimize distance errors between model and synthetic marker positions within the OpenSim musculoskeletal modeling software (Delp et al., 2007; Seth et al., 2018). For each of the three subjects studied, the JMP Tool employed a two-level optimization across eight shoulder motion tasks that included motions both with and without a handheld weight. The outer level optimization was performed in MATLAB (MathWorks, Natick, MA) and adjusted joint parameters and body scale factors, while the inner level optimization was performed by OpenSim (version 4.4) and involved repeated inverse kinematics analyses using the current guess for joint parameters and body scale factors. To evaluate model generalizability, we conducted a leave-one-out cross validation analysis for three subjects.

**Figure 1.**
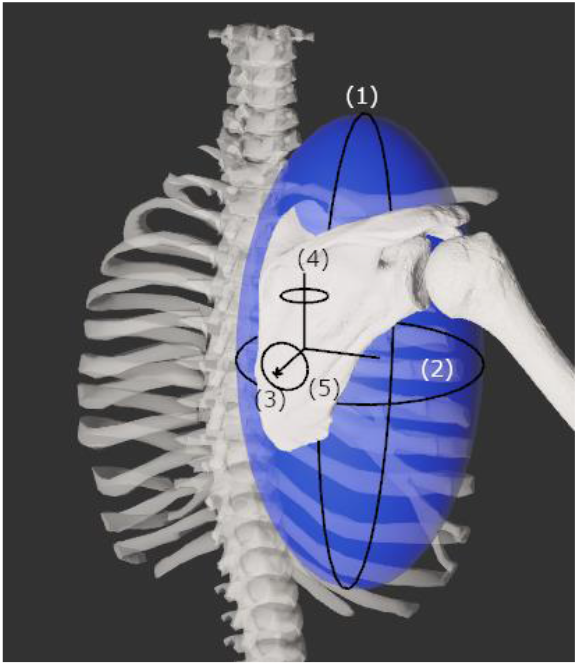
Diagram showing the five degrees of freedom of the scapula. The first two degrees of freedom are spherical coordinate-like rotations about a torso-centered origin. The third degree of freedom is a translation away from the torso-centered origin. The fourth and fifth degrees of freedom are rotations about the scapula-centered origin.

### 2.2 Experimental Data Preparation

To support the personalization and evaluation of the different shoulder models, we generated OpenSim-compatible subject-specific geometry and synthetic marker data for three subjects whose data were available in an open-source repository (Aliaj et al., 2021; Henninger, n.d.). This repository includes subject-specific polygonal geometry models (scapula and humerus) created from the CT images and biplane fluoroscopy kinematic data (100 Hz) for the scapula and humerus for a cohort of 20 healthy right-hand dominant adults separated by age and gender, where none of the subjects had any history of shoulder pathology (Kolz et al., 2021). The first subject was a male over the age of 45 (subject O45_002), the second subject was a female over the age of 45 (subject O45_001), and the third subject was a male under the age of 35 (subject U35_004).

We converted the subject-specific repository data into OpenSim-compatible subject-specific geometry and synthetic shoulder marker data in two steps. The first step involved conversion of subject-specific geometry models into OpenSim-compatible models. The coordinate systems in the repository geometry were modified using Geomagic Wrap (3D Systems, Morrisville, NC) so that they matched ISB anatomical axis definitions (Wu et al., 2005). The scapular and humeral polygonal geometry were then added to OpenSim Body objects in a generic OpenSim model, where the two bodies were connected to ground via 6 DOF joints. Virtual surface markers were placed on the scapula and humerus of the OpenSim Body objects using anatomical landmarks provided in the repository. For the scapula, these landmarks were the glenoid center (GC), the inferior angle (IA), the trigonum spinae (TS), the posterolateral acromion (PLA), and the acromioclavicular joint (AC), while for the humerus, they were the humeral head center (HHC), the elbow lateral epicondyle (LE), and the elbow medial epicondyle (ME).

The second step involved converting the 6 DOF motions of the scapula and humerus measured using biplane fluoroscopy into corresponding synthetic marker motions for those two bodies. This step was needed since the JMP tool uses surface marker data and repeated inverse kinematics analyses to find optimal functional axis locations and orientations in the body segments. To evaluate how accurately the shoulder model could reproduce experimental motions across different Cardinal planes, we selected a set of eight shoulder activities. These activities included a combination of weighted and unweighted overhead tasks during forward elevation, scapular plane abduction, coronal plane abduction, and internal/external humeral rotation. A custom MATLAB script converted the quaternion-based kinematics of the scapula and humerus obtained from the repository into Euler angle-based kinematics usable in OpenSim. Another custom MATLAB script used OpenSim functionality to generate synthetic marker data from the 6 DOF kinematics of the scapula and humerus. For each time frame across each activity, the scapula and humerus were placed in the poses defined by their experimental kinematic data and the three-dimensional position of each anatomical landmark in the ground coordinate system was recorded as a “synthetic marker” position. Because the experimental kinematic data were provided with respect to the torso coordinate system, three additional time-independent static markers (incisura jugularis; IJ, 7th cervical vertebra; C7, 8th thoracic vertebra; T8) were added to represent anatomically placed torso markers and allow for personalization of a shoulder model that included a generic torso segment. Thus, synthetic marker trajectories for each activity were generated for a total of 11 markers: five scapula, three humerus, and three torso. The synthetic marker positions were written to an OpenSim marker (.trc) file for each of the eight motions, and these marker positions served as gold-standard kinematics when personalizing each subject’s shoulder model using the JMP Tool.

### 2.3 Computational Tool Description

For the present study, the JMP tool within the Neuromusculoskeletal Modeling Pipeline software package (https://simtk.org/projects/nmsm) was used to personalize different shoulder kinematic models. The JMP tool is an easy-to-use tool that can personalize joint parameters, body scale factors, and marker positions in OpenSim models so as to minimize inverse kinematics marker distance errors. Every joint in an OpenSim model has six joint parameters (three translational, three rotational) for both the parent and child reference frames to define the joint’s transformational offsets. Each of these 12 joint parameters (six parent and six child) can be selected to be optimized using the JMP tool.

The goal of optimizing the joint parameters is to minimize the sum of squares of marker distance errors between experimental and virtual marker positions after the model’s pose has been optimized by the OpenSim Inverse Kinematics tool (Delp et al., 2007; Seth et al., 2018). The JMP tool accepts an XML settings file, a scaled generic model, and one or more .trc files for personalization. The JMP tools allows for multiple ‘tasks’ such that each task can accept a single marker file and a number of tool settings, indicating which joint parameter, body scale factor, and marker position design variables should be modified to minimize marker distance errors via inverse kinematics. If a user wants to personalize design variables for multiple motions, the motions of interest can be concatenated into a single, long marker (.trc) file.

### 2.4 Kinematic Model Formulation

For this study, we represented the shoulder complex by formulating two separate kinematic models: the first was an open-chain kinematic model of only the scapula independent of the clavicle and humerus, while the second was a closed-chain kinematic model of the shoulder complex. The open-chain scapula-only models were investigated using experimental data from only the first subject (male, over 45 years old), whereas the closed-chain shoulder complex models were investigated using experimental data from all three subjects. Experimental data from three subjects were used for the closed-chain shoulder complex model because additional analysis was conducted.

Three versions of the open-chain scapula-only kinematic model were implemented based on a previous study by Maurel et al. (1996) (Maurel et al., 2002). The three versions differed in the number of scapular DOFs (3, 4 or 5) with respect to the thorax. The first model possessed three DOFs consisting of two orthogonal thorax-centered spherical rotations followed by a translation to allow for scapular motion along an assumed ellipsoidal surface (i.e., spherical coordinates). The second model possessed four DOFs consisting of the three DOFs from the first model plus a fourth DOF to allow scapula-centered rotation about an approximately medial-lateral axis, representing sagittal plane rotations of the scapula with respect to the thorax. The third model possessed the four DOFs of the second model plus a fifth DOF to allow scapula-centered rotation about an approximately anterior-posterior axis, representing frontal plane rotations of the scapula with respect to the thorax. Using the MATLAB OpenSim application programming interface (API), we created all three scapula models using massless intermediate reference frames to allow for the correct sequence of coordinate motions without breaking compatibility with the JMP Tool.

Subsequently, two versions of the closed-chain shoulder complex kinematic model were implemented based on assessment of the three open-chain scapula-only kinematic models. The two closed-chain versions expanded the best two scapula-only models (4 and 5 DOFs) into closed-chain models that included the clavicle and humerus (details in section 2.6 Kinematic Model Evaluation). Accounting for DOFs eliminated by closed-chain constraints, the closed-chain shoulder model incorporating the 4 DOF open-chain scapula model possessed 6 independent DOFs (3 for the scapula and clavicle + 3 for the humerus), while the closed-chain shoulder model incorporating the 5 DOF open-chain scapula model possessed 7 independent DOFs (4 for the scapula and clavicle + 3 for the humerus).

The two closed-chain models were constructed in five steps. First, a generic torso body (6 DOF) was added to an empty OpenSim model. Second, a subject-specific scapula body (either 4 DOF or 5 DOF) was added as a child body of the torso. Third, a subject-specific humerus body (3 DOF) was added as a child body of the scapula using ISB standard coordinate system definitions for internal rotation, elevation, and elevation plane (Wu et al., 2005; Seth et al., 2016). Fourth, a scaled generic clavicle body (2 DOF) was added as a child body of the torso. Fifth, a point constraint was added between the AC landmark on the scapula and the end of the clavicle to close the kinematic chain.

### 2.5 Kinematic Model Personalization & Evaluation

Using the JMP tool and the proposed scapula-only and kinematic models, we developed a repeatable process to personalize both the open-chain scapula-only kinematic models and the closed-chain shoulder complex kinematic models. To generate the input marker data needed by the JMP tool, we concatenated synthetic marker data (100 Hz) from all eight experimental motions into a single marker file (.trc). For each open-chain scapula-only model, a single JMP task was created to personalize all relevant parameters in the scapulothoracic joint. In contrast, for each closed-chain shoulder complex model, a two-stage personalization process was used. First, a JMP task was created to personalize each anatomical joint (e.g., glenohumeral, acromioclavicular, sternoclavicular, and scapulothoracic) separately in a specific order to minimize the risk of finding a local minimum during the personalization process. Second, a final JMP task was created to personalize all anatomical joints together.

To personalize the scapulothoracic joint in each scapula-only model, we created a single JMP task within a single JMP XML settings file that personalized relevant scapulothoracic joint parameters. Specifically, the settings file allowed the JMP tool to adjust the orientation of the parent frame fixed in the torso body, the location of the parent frame fixed in the torso, and the location and orientation of the child frame fixed in the scapula body.

To personalize each joint separately in each shoulder complex model, we created a sequence of five JMP tasks within a single JMP XML settings file that personalized relevant joint parameters and/or body scale factors systematically for one anatomical joint at a time. For these five tasks, the concatenated synthetic marker data were downsampled to 20 Hz to improve computational speed. The first JMP task focused on the glenohumeral joint and adjusted the location of the glenohumeral joint center in both the scapula and humerus bodies. The second JMP task focused on the acromioclavicular joint and adjusted the scaling of the generic clavicle body and the location of the point constraint in the clavicle body. The third JMP task focused on the sternoclavicular joint and adjusted the orientation of the parent frame fixed in the torso body. The fourth JMP task focused on the scapulothoracic joint and adjusted the location of the parent frame fixed in the torso body. The fifth JMP task also focused on the scapulothoracic joint and adjusted the location and orientation of the child frame fixed in the scapula body.

To personalize all joints together in each closed-chain shoulder complex model, we created a final JMP task within a single JMP XML settings file that personalized all joint parameters and body scale factors described above for all anatomical joints together. The full sample of concatenated synthetic marker data (100hz) were used for this task.

We quantified the ability of each personalized kinematic model to reproduce the gold-standard synthetic marker motion data in two ways. First, we personalized each of the open-chain scapula-only models (3, 4, and 5 DOF) to assess how well each model could reproduce the gold-standard marker motion data. Experimental data from only the first subject (male, over 45 years old) were used to evaluate the different open-chain scapula-only models, and marker distance errors were calculated using only markers placed on the scapula. We defined an average and maximum accuracy threshold (avg error = 5.0 mm | max error = 15.0 mm) that each scapula-only model needed to reach to be included in a closed-chain shoulder complex model. Once evaluation of the scapula-only models was completed, we personalized the closed-chain shoulder complex model (scapula, clavicle, and humerus) using the scapula-only models that fell below the specified average accuracy threshold. Experimental data from all three subjects were used to evaluate the different closed-chain shoulder complex models, and marker distance errors were calculated using markers placed on the scapula, torso, and humerus.

To evaluate the kinematic accuracy of each personalized open-chain scapula only and closed-chain shoulder complex model, we calculated average and maximum marker distance errors (always positive) for each marker used. Marker distance errors were calculated by leveraging the JMP tool’s plotting functionality. Average marker distance errors indicate the model’s ability to match poses that the subject moves through frequently, while maximum marker distance errors indicate the model’s ability to match the most challenging pose.

In addition, for the most accurate closed-chain shoulder complex model identified, we performed a leave-one-out cross validation analysis to quantify the model’s ability to match kinematics not included in the personalization process. To perform the cross-validation analysis, we personalized each subject’s best closed-chain model using concatenated gold-standard data from seven motions. We then calculated marker distance errors using gold-standard data from the eighth motion. This process was repeated eight times so that each of the eight experimental motions was left out once from the personalization process and used for testing purposes. For each omitted motion, average marker distance errors between model and gold-standard marker positions were calculated using OpenSim inverse kinematics analyses. We calculated a final average marker distance error representing the average error across all omitted motions.

## 3 Results

The kinematic errors produced by the open-chain scapula-only models decreased with increasing number of scapula DOFs (Figure 2). The 3 DOF scapula-only model showed higher marker distance errors (avg error = 5.7 mm | max error = 20.9 mm) compared to the 4 DOF (avg error = 2.8 mm | max error = 12.5 mm) and 5 DOF (avg error = 0.8 mm | max error = 5.2 mm) models. Since only the 4 and 5 DOFs models met the average and maximum accuracy threshold of 5.0 mm and 15.0 mm, respectively, only those two models were incorporated into subsequent closed-chain shoulder complex models.

**Figure 2.**
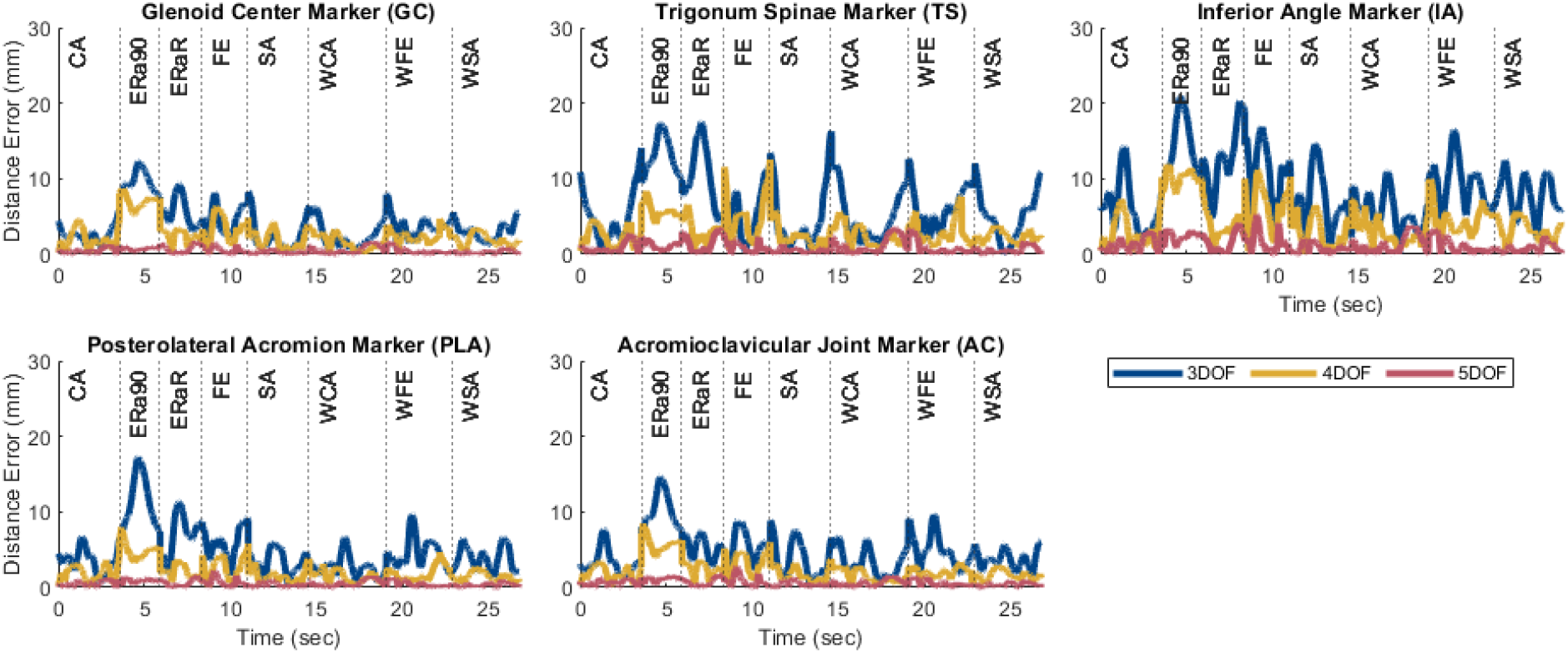
Plots showing the marker distance error of each marker throughout the eight motion tasks after personalization for the 3, 4, and 5 DOF scapula-only models. The motions included a combination of weighted (prepended with ‘W’) and unweighted overhead tasks during forward elevation (FE, WFE), scapular plane abduction (SA, WSA), coronal plane abduction (CA, WCA), and internal/external humeral rotation (ERa90, ERaR).

After the closed-chain shoulder complex models that incorporated 4 and 5 DOF scapula models were personalized, the kinematic errors produced by the closed-chain models again decreased with increasing number of scapula DOFs (Figure 3).

**Figure 3.**
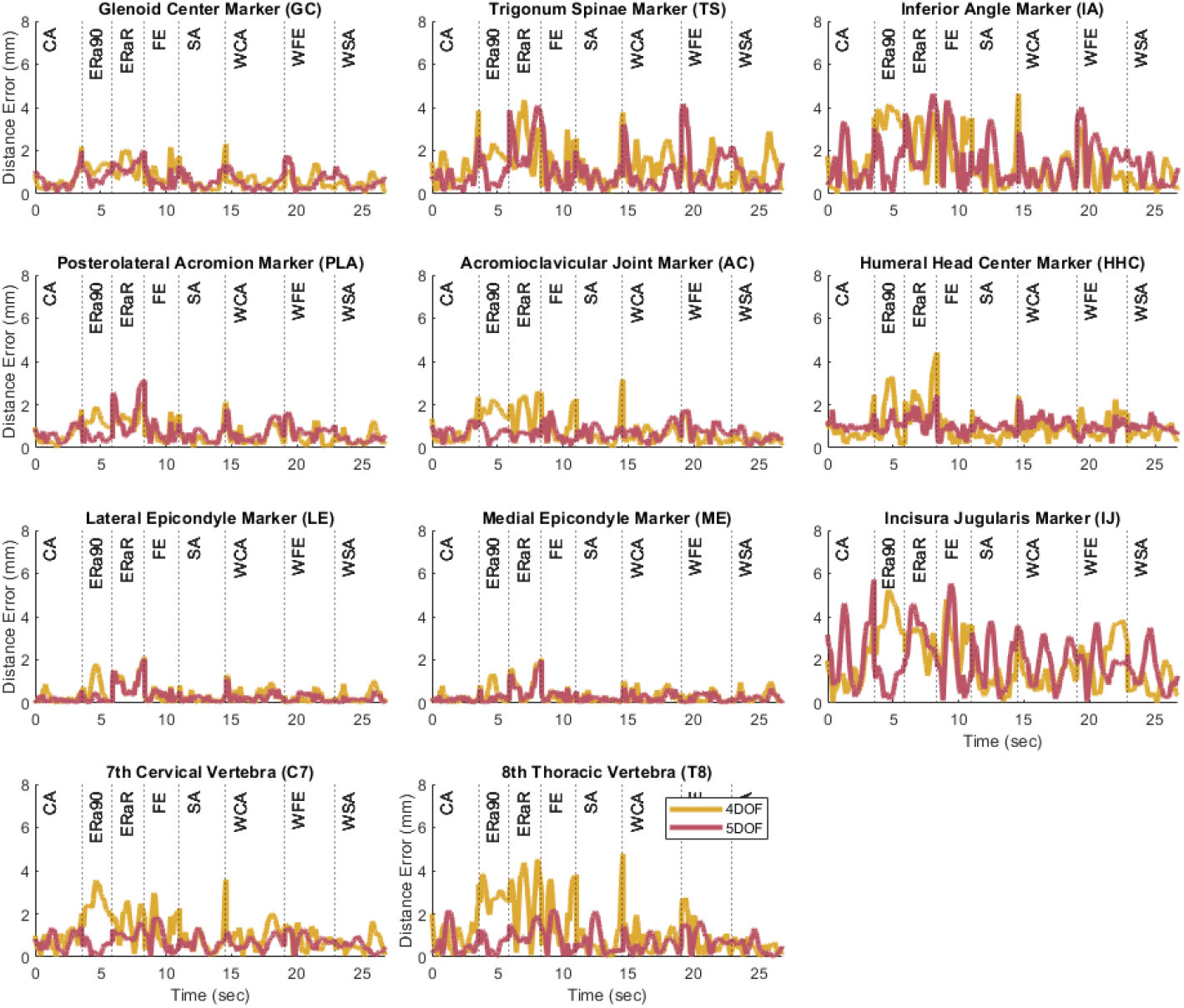
Plots showing the marker distance error of each marker throughout the eight motion tasks after personalization for the 4 and 5 DOF closed-chain shoulder complex models.

The closed-chain shoulder model with a 4 DOF scapula showed higher marker distance errors (avg error = 1.0 mm | max error = 5.2 mm) compared to the closed-chain shoulder model with 5 DOF scapula (avg error = 0.9 mm | max error = 5.7 mm). Since the closed-chain shoulder complex model with 5 DOF scapula had the lowest average marker distance errors (Figure 4), it was selected for use in the subsequent leave-one-out cross validation analysis.

**Figure 4.**
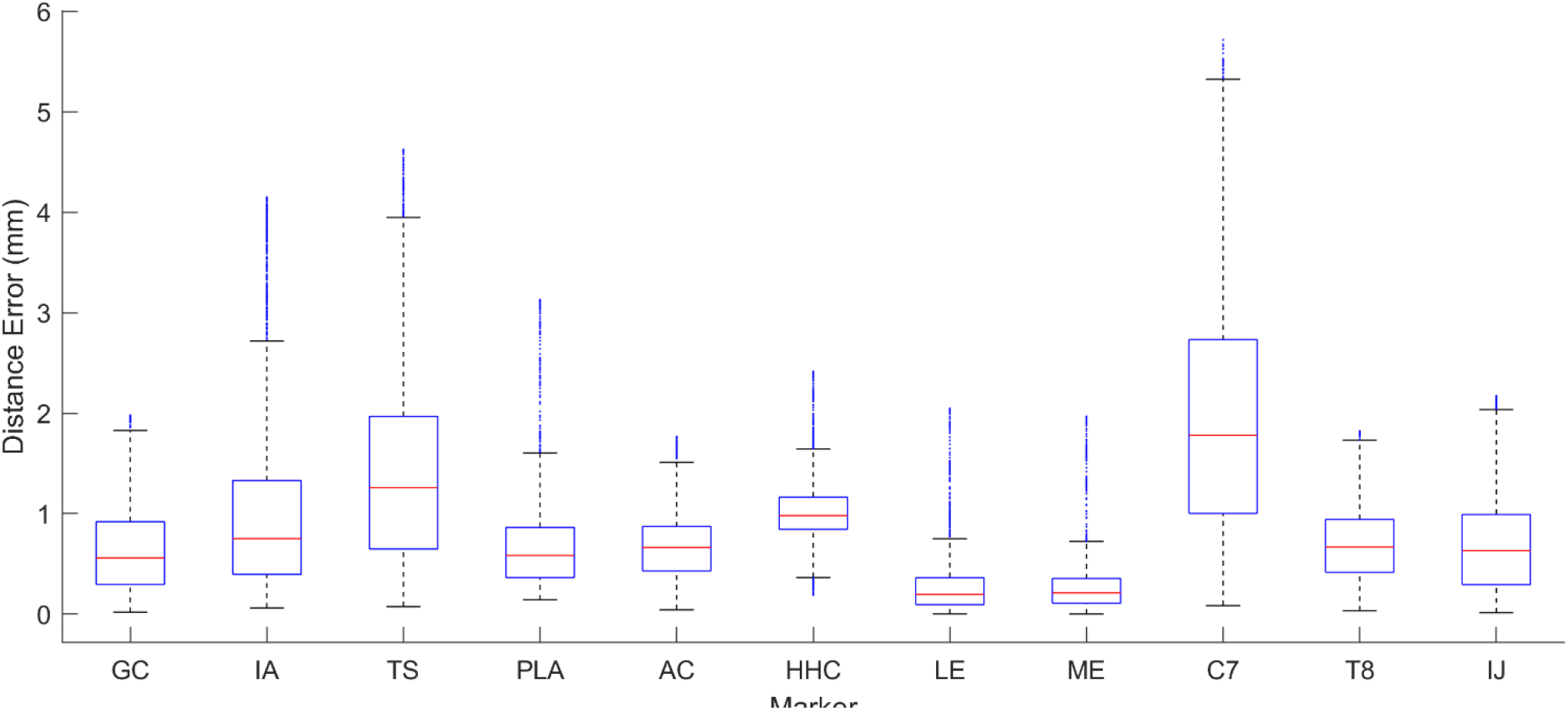
Box plot showing the range of distance errors across the eight motion tasks after personalization for the 5 DOF closed-chain shoulder complex model.

The cross validation analysis showed similar average omitted motion marker distance errors across all three subjects (Table 1). For each personalized closed-chain shoulder complex model incorporating a 5 DOF scapula, average omitted motion marker distance errors were 1.0 mm, 1.3 mm, 1.4 mm for subjects 1, 2, and 3, respectively. These errors were only slightly larger than the corresponding errors found when each model was personalized using gold-standard data from all eight motions together.

**Table 1.**
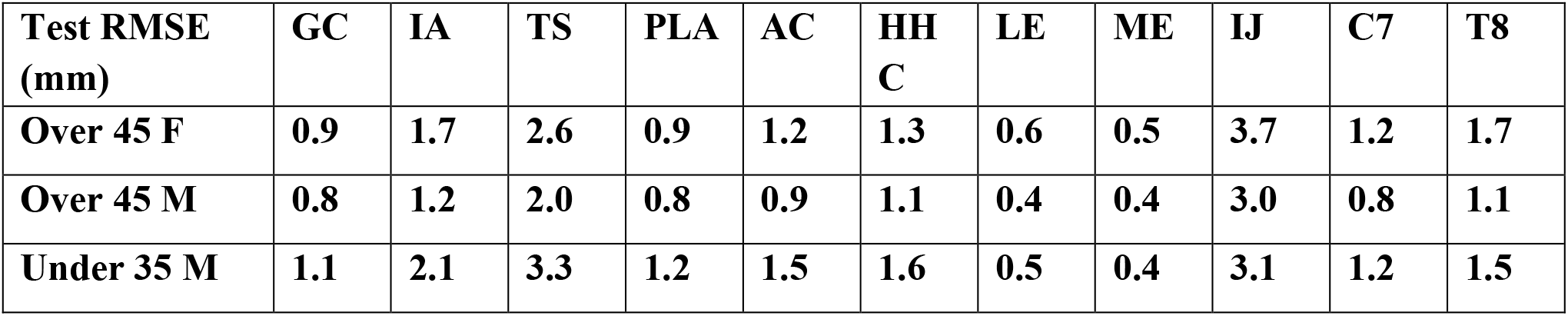
The average marker distance error across all test motions in the leave-one-out cross validation analysis.

## 4 Discussion

The objective of this work was to develop a novel methodology for creating accurate personalized shoulder kinematic models. Using subject-specific kinematic data from bi-plane fluoroscopy and associated subject-specific scapula and humerus geometry, we developed a robust personalization process using the JMP tool. We demonstrated that personalized open-chain scapula-only models possessing either 4 or 5 DOFs relative to the thorax achieved acceptable kinematic accuracy relative to gold standard experimental data. We also demonstrated that a personalized closed-chain shoulder complex model (scapula+clavicle+humerus+thorax) possessing 7 independent DOFs and incorporating a 5 DOF scapula model produced the most accurate tracking of gold-standard kinematic data with an average marker distance error of 0.9 mm across eight motions. Based on the results of a cross validation analysis performed for three healthy subjects of different ages and sexes using the 7 independent DOF closed-chain shoulder model, the kinematic model personalization methodology developed using the JMP tool seems to be robust for healthy shoulders.

Several interesting results were found during the investigation of this novel model personalization process. During the open-chain, scapula-only model personalization, the three models (3, 4, and 5 DOF) increased in accuracy as the number of degrees of freedom increased. This outcome was likely due to the increased DOFs allowing for more complex scapular motion. Once the scapula-only models were added to the closed-chain shoulder complex model, the 4 DOF model introduced significant errors at the ends of the motion trials, especially for movement tasks with a high range of motion. The closed-chain shoulder complex model with the 5 DOF scapula showed the lowest average marker distance errors, but not the lowest maximum marker distance error across all the tasks simulated. Despite the maximum marker distance error of the 5 DOF scapula model being 0.5 mm greater than the 4 DOF scapula model, the 5 DOF scapula model had a lower final cost upon completion of the JMP tool (5 DOF: 1.38, 4 DOF: 1.76). This result indicates that the JMP tool was able to find a more accurate solution with the 5 DOF model. Since the JMP tool’s cost function minimizes the sum of the squares of the marker distance errors, explicit reduction of maximum values is not possible with JMP and adding a feature to reduce maximum values would be difficult since a maximum error value would not be differentiable. The cross-validation analysis produced comparable marker distance errors (range: 1.0 mm - 1.4 mm) across all three subjects, demonstrating that this model structure is likely generalizable across different age and sex groups, at least for healthy shoulders.

Our novel kinematic model personalization framework differs from previous shoulder modeling methods to facilitate high accuracy and kinematic personalization. Rather than using a closed-chain kinematic model, previous shoulder modeling studies have attempted to model the scapulothoracic joint using either a mobilizer to add a degree of freedom to the scapula or through regression-based modeling of the clavicle and scapula relative to the position of the humerus (Holzbaur et al., 2005; Seth et al., 2016; Wu et al., 2016). Our approach leverages the JMP tool to facilitate both ellipsoidal-like and body-centered rotations of the scapula body inside a closed-chain shoulder model.

Development of a closed-chain shoulder model that moves realistically and matches experimental data is difficult using previously published methods, but with the JMP tool, we have created highly accurate subject-specific closed-chain shoulder models. These personalized closed-chain shoulder models lead to more consistent and realistic motions of the scapula, which is critical for future applications involving motion prediction and treatment optimization.

To facilitate development of a highly accurate kinematic model of the scapula, we made two assumptions about how to define the pose of the scapula relative to the thorax that differ from previous shoulder models. First, unlike the model developed by Saul et al., we selected scapula generalized coordinates that did not match clinical definitions of shoulder motion (Holzbaur et al., 2005). Freeing ourselves from this limitation allowed us to define scapular generalized coordinates whichever way made the most physical sense. Second, unlike the model developed by Seth et al., we did not constrain scapular motion to remain in contact with an ellipsoidal surface (Seth et al., 2016; Blache et al., 2024). Instead, we defined the first three scapular DOFs using spherical coordinates, which allowed the translation to change with the two rotations. The drawback of including a translational generalized coordinate is that motion along this translational coordinate is influenced by not only muscle forces but also scapulothoracic contact forces, complicating subsequent dynamic modeling of scapular motion.

One way to eliminate this issue is to model scapular translation as a function of the first two scapular rotations. To explore how well this approach could work, we modified our 7 DOF closed-chain shoulder complex model by adding an OpenSim Coordinate Coupler constraint to make scapular translation a linear function of the first two scapular rotations. A linear function was chosen since OpenSim does not currently support ellipsoidal functions. For each subject, the coefficients of the linear function were obtained by regression fitting results from the subject’s 7 DOF closed-chain model without the coordinate coupler, which yielded an average absolute fitting error of 1.4 mm across all three subjects. When this linear function was implemented as a coordinate coupler constraint for each subject, the modified 6 DOF closed-chain model was able to match gold standard marker position data with average marker distance errors of 2.9 ± 1.0 mm, 3.2 ± 1.5 mm, 3.1 ± 1.4 mm for subjects 1, 2, and 3, respectively. Though these average errors are roughly three times those obtained without the coordinate coupler constraint, these errors are still quite low and could be improved if a different functional relationship (e.g., ellipsoidal) could be used. Future work will explore implementing a method for using any general functional relationship for an OpenSim coordinate coupler, as this approach could facilitate a more accurate shoulder kinematic model with fewer DOFs while eliminating the need for a contact model to help control scapular translation during dynamic simulations.

Two features of the JMP tool are particularly important for the development of personalized shoulder kinematic models. First, during a closed-chain inverse kinematic calculation, if the model is unable to assemble the bodies (i.e., “close the loop”) due to inconsistent design variable values, the tool returns a large error for that time frame so that the optimization will continue and will avoid that region of the solution space. Second, when a point constraint is included in an OpenSim model, if a body associated with the point constraint is scaled, the point constraint remains at its original location. However, the JMP tool also provides the option for a point constraint location to be updated when the associated bodies are scaled. For the purposes of this study, the point constraint location was allowed to scale along with the clavicle body.

This study possesses at least three important limitations that are worth highlighting. First, only gold-standard synthetic marker data was used. Although this choice allowed us to isolate the accuracy of the kinematic model from the accuracy of the experimental data, it is not known how the accuracy and personalization process will change to best accommodate skin-based marker data. Second, only three subjects from a dataset of healthy individuals were used to develop our models. Other healthy or pathological datasets may not produce similar kinematic accuracy. Third, the personalization protocol used in this study was not the result of an exhaustive investigation of all possible personalization protocols. Best practices related to the use of the JMP tool have not yet been explored, and a better personalization protocol could potentially be found through further investigation. Despite these limitations, we developed a novel approach to create a highly accurate, closed-chain kinematic shoulder model that can reproduce 8 separate motion tasks to within 1.4 mm of average error.

In conclusion, this study developed personalized shoulder models of varying accuracy, complexity, and constraint using the JMP Tool within the Neuromusculoskeletal Modeling Pipeline software (Hammond et al., 2024). The tool can facilitate high quality marker-based personalized kinematic modeling of the shoulder and will likely be equally valuable when applied to other joints as well. Using a musculoskeletal model that achieves high kinematic accuracy is important because errors in joint locations and orientations lead to errors in inverse kinematics results, which in turn lead to errors in inverse dynamics results, which in turn lead to errors in EMG-driven model calibration results, which ultimately lead to errors in predictive simulation results (Reinbolt et al., 2007). The shoulder kinematic models developed here can be used as the foundation for a future shoulder model that includes personalized muscles as well as all of the features required for solving torque and synergy-driven direct collocation optimal control problems. Future work will explore how well the same shoulder model personalization process works using experimental motion capture data collected from markers placed on the skin (Seth et al., 2016; Maier et al., 2024).

## 5 Data Availability Statement

The experimental source data used in this study can be found at 10.5281/zenodo.4536683 (newly released database can be found at 10.5281/zenodo.10972004), while the modeling data used in this study can be found at https://simtk.org/projects/shoulder-model

## 6 Conflict of Interest

The authors declare that the research was conducted in the absence of any commercial or financial relationships that could be construed as a potential conflict of interest.

## 7 Author Contributions

HH provided the experimental data. CH performed all model personalization tasks, prepared the figures, and drafted the manuscript. CH, BF, and JG analyzed the data. CH, BF, and JG interpreted the results of analyses. All authors revised the manuscript.

## 8 Funding

This work was conducted with support from the Cancer Prevention and Research Institute of Texas (grant RR170026 to BF), the National Institutes of Health (R01 EB030520 to BF), the National Science Foundation (1404767, 1159735, 1052754, and 1842494) and a collaborative exchange grant through the Orthopaedic Research Society (to JG). Funding from the National Institutes of Health (R01 AR067196 to HH) was used to generate the original experimental data.

## Notes

https://zenodo.org/records/10972005

https://simtk.org/projects/shoulder-model

### Competing Interest Statement

The authors have declared no competing interest.

